# Human Brain Aging is Associated with Dysregulation of Cell-Type Epigenetic Identity

**DOI:** 10.1101/2024.06.01.596981

**Authors:** Hyeonsoo Jeong, Isabel Mendizabal, Soojin V. Yi

## Abstract

Significant links between aging and DNA methylation are emerging from recent studies. On the one hand, DNA methylation undergoes changes with age, a process termed as epigenetic drift. On the other hand, DNA methylation serves as a readily accessible and accurate biomarker for aging. A key missing piece of information, however, is the molecular mechanisms underlying these processes, and how they are related, if any. Addressing the limitations of previous research due to the limited number of investigated CpGs and the heterogeneous nature of tissue samples, here we have examined DNA methylation of over 20 million CpGs across a broad age span in neurons and non-neuronal cells, primarily oligodendrocytes. We show that aging is a primary predictor of DNA methylation variation, surpassing the influence of factors such as sex and schizophrenia diagnosis, among others. On the genome-wide scale, epigenetic drift manifests as significant yet subtle trends that are influenced by the methylation level of individual CpGs. We reveal that CpGs that are highly differentiated between cell types are especially prone to age-associated DNA methylation alterations, leading to the divergence of epigenetic cell type identities as individuals age. On the other hand, CpGs that are included in commonly used epigenetic clocks tend to be those sites that are not highly cell type differentiated. Therefore, dysregulation of epigenetic cell-type identities and current DNA epigenetic clocks represent distinct features of age-associated DNA methylation alterations.

## INTRODUCTION

Aging has a profound influence on DNA methylation. Studies dating back several decades have demonstrated DNA methylation levels of specific CpGs exhibiting age-associated changes (e.g., [1–3]. Technical advances in the last decade led to the development of relatively cost-friendly microarray methods to study DNA methylation of many CpGs. Subsequently, DNA methylation profiling of large cohorts and from different tissue types followed (e.g., [4–9]). To study the effect of aging on DNA methylation, these studies typically performed either correlation and/or linear regression analyses between DNA methylation and age, identifying numerous CpGs that showed significant variation with aging. The results of these studies solidified that aging has fundamental impacts on DNA methylation.

It was shown that DNA methylation of genetically identical monozygotic twins also diverge with aging [10], indicating that age-associated DNA methylation changes are not necessarily programmed in the genome (however, see [11] for a different interpretation). A term ‘epigenetic drift’ is often used to refer to changes of DNA methylation that occur during aging [12–14]. While there was some earlier disagreement over the nature of epigenetic drift regarding whether it involves decrease or increase of DNA methylation, it became apparent that both patterns were prevalent. Promoters and CpG islands, which tend to have lower levels of DNA methylation (e.g., [15–17]), often undergo hypermethylation with aging, while intergenic/repetitive regions with higher DNA methylation tend to experience hypomethylation [14, 18, 19]. These patterns support the idea that epigenetic drift might be due to gradual dysregulation of epigenetic maintenance over the lifespan, a pattern we will demonstrate more clearly in this work using nucleotide-resolution data of nearly all CpGs in the human genome.

A landmark in age-associated DNA methylation research is the development and application of the so-called ‘DNA methylation clocks’ [9, 20–24]. Specifically, DNA methylation of subsets of CpGs can be used as predictors of age. They are often identified using supervised machine learning methods applied to large cohorts, constructed using data from single tissue or multiple tissues [21, 23]. DNA methylation clocks are remarkably robust, and in many cases perform better than other traditional predictors of biological aging [21]. Epigenetic age predictors have wide-ranging applicability for the study of human health and medicine.

These two aspects of age-associated DNA methylation changes, namely epigenetic drift and DNA methylation clocks, open many questions and opportunities to study aging from the perspective of epigenetic programs over lifespan [18, 21, 23]. However, there are several current deficiencies of knowledge that are critical to fully understanding and utilizing these patterns. For example, is epigenetic drift a feature of all CpGs in the genome? Previous studies typically used subsets of CpGs, often relying on DNA methylation arrays which were biased toward promoters that are not necessarily variable [25]. Therefore, investigations utilizing unbiased, whole-genome CpGs are necessary to provide an objective, genome-wide view. Second, how different or similar are epigenetic changes in distinct tissues and cell types? Many previous studies used bloods, due to the ease of sampling. However, even though blood sampling remains as a powerful tool to investigate aging, data from distinctive cell types can provide additional insights into the mechanistic, cell-type resolved aspects of aging. Third, what is the relationship between epigenetic clocks and epigenetic drift? Fundamentally, how do the epigenetic drift and epigenetic clocks relate to the underlying biological mechanisms of aging?

To address these questions, we need to extend the study of age-associated DNA methylation changes to the whole genome, using methods that can interrogate all genomic CpGs, such as the whole-genome bisulfite sequencing (WGBS). Another key missing piece of information in addressing these issues is understanding DNA methylation at cellular resolution [21, 23]. As epigenomic studies begin to reveal substantial heterogeneity of cellular epigenetic programs, it is necessary to evaluate how age-associated DNA methylation changes occur in different cell populations. Unfortunately, obtaining genome-wide single-cell profiles of DNA methylation is technically challenging and cost prohibitive for most samples, with current technologies. An effective approach is through the use of whole-genome bisulfite sequencing on purified cell-types. Here, we present our analyses of extensive whole-genome bisulfite sequencing data sets of DNA methylation from neurons and non-neuronal cells (primarily oligodendrocytes), separated by fluorescence-activated nuclei sorting, from individuals across a broad age span (ranging from neonate to 85 years). We demonstrate the impact of epigenetic drift across the whole genome and identified CpGs that show significant DNA methylation changes with aging, using a regression method specifically developed for the analysis of WGBS data. Our work uncovers a novel property of age-associated changes in cell-type specific epigenomes, and provides a list of top candidate positions that undergo age-associated changes in two major cell types in the human brain, namely neurons and oligodendrocytes.

## RESULTS

### Age is a Major Driver of DNA Methylation Change in the Whole Genome CpGs

To gain insight into the aging programs in the human brain at cell type resolution, we examined 127 whole-genome bisulfite sequencing (WGBS) data sets of neurons (NeuN+, N=77), oligodendrocytes (OLIG2+, N = 42) and non-neuronal cells (NeuN-, N = 8) from the dorsolateral prefrontal cortex, collected from two independent studies [26, 27] (Supplementary Table 1). These data were from individuals across a broad age span, ranging from 2.4 months to 85 years (mean = 43.5 years, Supplementary Table 1). To avoid erroneous methylation calls due to genetic polymorphisms at cytosine bases, we first mapped all the matched whole genome sequencing data and excluded positions that were polymorphic at cytosines (Methods). Consequently, we were able to determine DNA methylation levels of 23.6 million CpGs from these data with a minimum coverage of 5x (over 85% of all CpGs in the human genome). Hierarchical clustering analysis indicated that NeuN+ samples from the two data sets clustered together, while OLIG2+ and NeuN-clustered together (Figure 1A).

**Figure 1.**
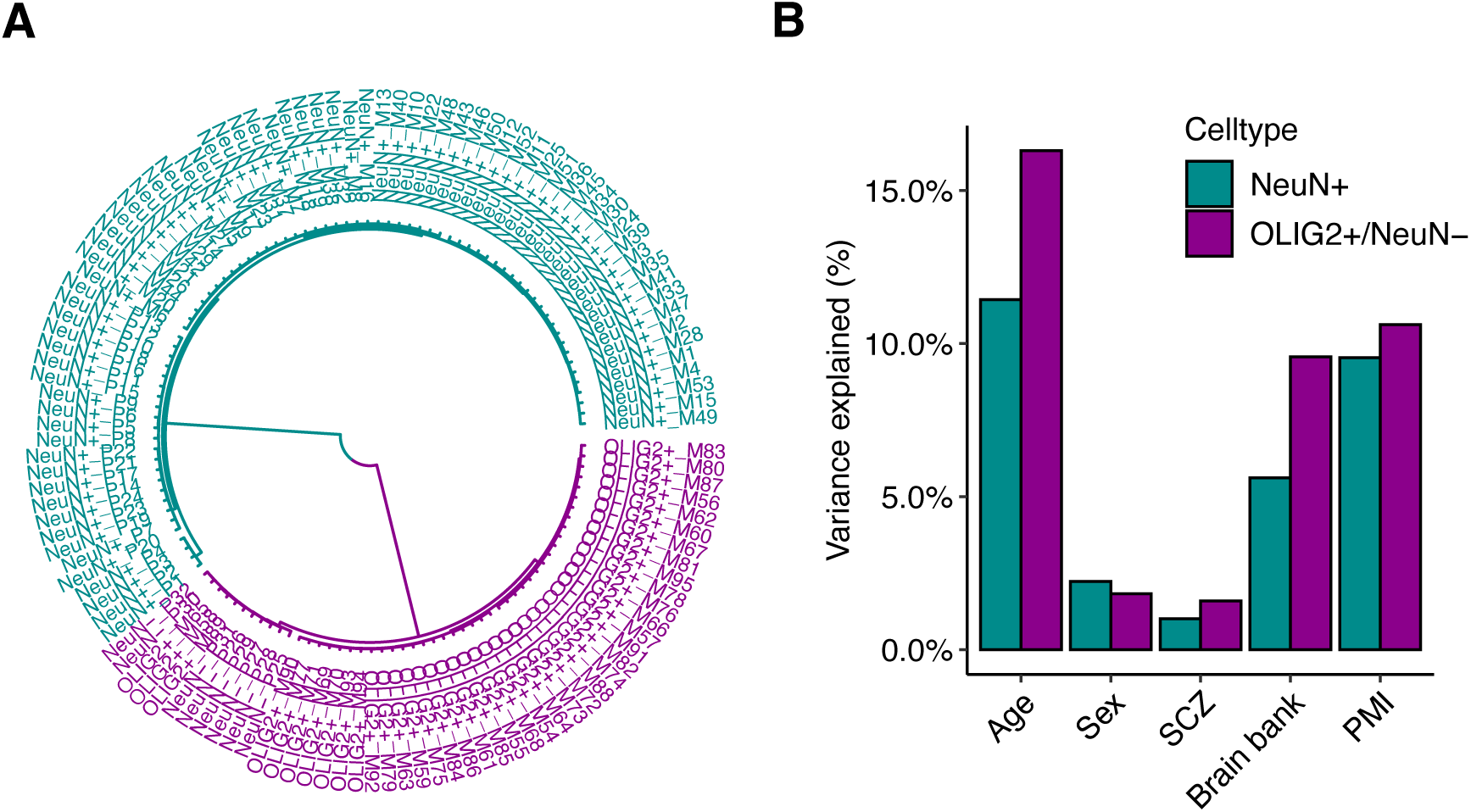
Variation of DNA methylation in neurons and oligodendrocytes/non-neuronal cells. A) Hierarchical clustering of 127 whole-genome bisulfite sequencing (WGBS) data used in this study separated two clusters of cell populations (NeuN+ and OLIG2+/NeuN-). B) Proportion of variation explained by each variable after controlling for the effect of genetic contribution (SCZ, schizophrenia; PMI, post-mortem interval).

Age was a main determinant of variation of DNA methylation in the analysis of WGBS data sets. Overall, age explained a greater amount of variation of DNA methylation (11.4% and 16.3% for NeuN+ and OLIG2+, respectively) compared to other factors such as disease status (<1.6%) and sex (<2.2%, using CpGs in autosome) (Figure 1B). The tissue repository and postmortem interval of the samples also had impacts on DNA methylation. In the following analyses, we used these variables as covariates to assess the independent impacts of aging while accounting for these variables (Methods).

Also, it should be noted that samples from the Price et al. [27] were biased toward relatively young age groups (neonate to early 20s) while those from Mendizabal et al. [26] ranged from late 20s to 80s (Supplementary Table 1). Given that DNA methylation dramatically changed during early development [27, 28], DNA methylation change in Price et al. data set could be mainly driven by the effect of DNA methylation shift during early development. Principal components and pairwise correlation coefficients showed that DNA methylation levels were influenced by both the difference between early age and the two studies (shown for NeuN+, Supplementary Figure 1). Notably, DNA methylation in neonates was distinct from those of individuals from other ages (Supplementary Figure 1, [27]). In addition, the two data sets were also generated from two different labs, thus subject to slightly different protocols, sequencing platforms, and other potential yet unknown differences. Consequently, we treated the two data sets separately in the subsequent analyses, unless otherwise stated.

Previous studies have demonstrated that genome-wide DNA methylation changes with aging could be explained by the so-called epigenetic drift, one consequence of which was an increase of DNA methylation for sites that are initially lowly methylated, and a decrease of DNA methylation for highly methylated sites [14, 18, 19]. We used an average methylation level of neonates as a proxy for the putative ‘initial’ methylation state and asked if the direction of epigenetic drift was related to that. The relationship between these two variables represented significant genome-wide trends (Spearman’s rho = −0.32, and −0.12 respectively for neurons and oligodendrocytes, both *P* < 2.2X10^-16^, Supplementary Figure 2A). We observed the same trend when we use the mean methylation of individuals with ages less than 20 as a predictor (Supplementary Figure 2B). Nevertheless, the overall trends were noisy and relatively weak, highlighting many exceptions and variations of individual CpGs (see below).

In mammalian and other vertebrate genomes, DNA methylation levels are known to exhibit a clear bimodal distribution of lowly and highly methylated CpGs [16, 17]. Given the direction of epigenetic drift we have demonstrated above, it is expected that the genome-wide epigenetic drift diminishes differences between the extreme ends of DNA methylation (since heavily methylated CpGs lose DNA methylation and lowly methylated CpGs gain DNA methylation). Indeed, we observed that the clear bimodality was more dispersed in old adults, although the distributions vary across individuals (Supplementary Figure 3).

### Age-associated CpGs are Highly Cell-Type Specific

To characterize age-associated DNA methylation changes, we investigated age-associated DNA methylation changes using data from Mendizabal et al. [26]. This data set contained samples from the aging lifespan, rather than developmental shift. We applied a generalized linear model framework developed to analyze WGBS data [29]. Following these procedures, we identified 4,480 and 2,253 CpGs that showed significant age-associated methylation changes in NeuN+ and OLIG2+ (Supplementary Table 2 and 3, respectively). These CpGs were detected using the cutoff of FDR < 0.1, which corresponded to *P* < 3.18 x 10^-5^ and *P* < 1.52 x 10^-5^ for NeuN+ and OLIG2+, respectively. They were henceforth referred to as ‘age-differentially methylated loci’ or simply ‘age-DMLs’. We also performed the same analysis after combining the two data sets and found that test statistics from Mendizabal et al. data vs. the combined data sets were highly correlated (Spearman’s rho = 0.76) (Supplementary Figure 4). For example, we observed that 73% of age-DMLs identified from Mendizabal et al. data were overlapping with age-DMLs from the combined sets, indicating consistent patterns of age-associated DNA methylation changes in the two data sets.

Remarkably, age-DMLs of neurons and oligodendrocytes were highly distinct, with only 16 overlapping CpGs between them. Furthermore, DNA methylation changes with age showed distinct patterns between two cell types. In NeuN+, age-DMLs were dominated by those that increased DNA methylation with age (77.2%, also referred to as ‘age-hyper DMLs’). In contrast, age-DMLs in OLIG2+ were more evenly distributed, yet slightly biased toward hypomethylation with age (55.6%, also referred to as ‘age-hypo DMLs’) (Figure 2A). For example, in one of the most significant age-DML in NeuN+ (position 52,176,442 in Chromosome 3, intron 6 of 10 of POC1A gene), DNA methylation increased by 4% every decade for NeuN+, while in OLIG2+ that position remained constantly hypermethylated throughout the age span in the data (Figure 2B, left panel). In another example, an OLIG2+ specific age-DML in position 73,854,649 in Chromosome 3 (in intergenic region) showed a steady decline of DNA methylation throughout the age span in the data (Figure 2B, right panel).

**Figure 2.**
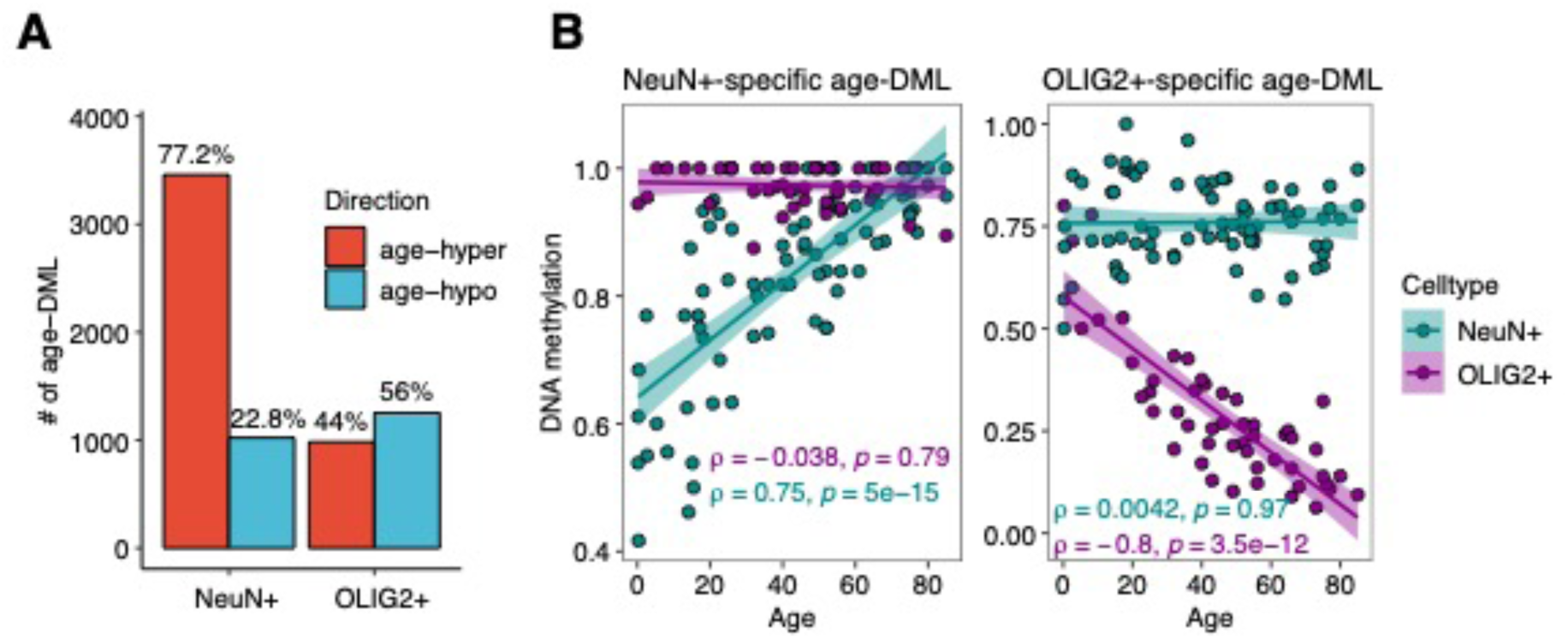
Age-associated methylation changes are highly cell-type-specific. A) Contrasting abundances of significant age-associated CpG methylation changes (age-DMLs) in NeuN+ and OLIG2+. B) Examples of NeuN+ or OLIG2+ specific age-DMLs. The first plot is an age-DML in NeuN+, located in an intron of POC1A gene. The correlation coefficients of NeuN+ and OLIG2+ are shown, demonstrating NeuN+ specific, significant increase of DNA methylation with age. The second plot is an age-DML in OLIG2+ located in intergenic regions of Chromosome 3, demonstrating a decrease of DNA methylation in OLIG2+ but not in NeuN+.

We have previously demonstrated that NeuN+ and OLIG2+ exhibiting highly distinctive genome-wide DNA methylation landscapes [26] and that the majority of these differences were evolutionarily conserved [30]. Remarkably, we found that age-DMLs were biased to cell-type specifically methylated CpGs. Age-DMLs were more than five-fold enriched in differentially methylated regions between neurons and oligodendrocytes (referred to as ‘cell-type DMRs’, [26]) (*P*<10^-5^, hypergeometric test, Figure 3A). CpGs whose DNA methylation increased with age in NeuN+ (NeuN+ age-hyper DMLs) were nearly exclusively (81.9%) the positions where NeuN+ was lowly methylated than OLIG2+, that is, decreasing cell-type differences in aging. Interestingly, OLIG2+ age-hypo DMLs were predominantly (78.5%) where OLIG2+ were less methylated compared to NeuN+, suggesting increased cell-type differences with age (Supplementary Figure 5). Globally, age-DMLs showed significantly greater cell type differences compared to randomly selected controls (Figure 3B).

**Figure 3.**
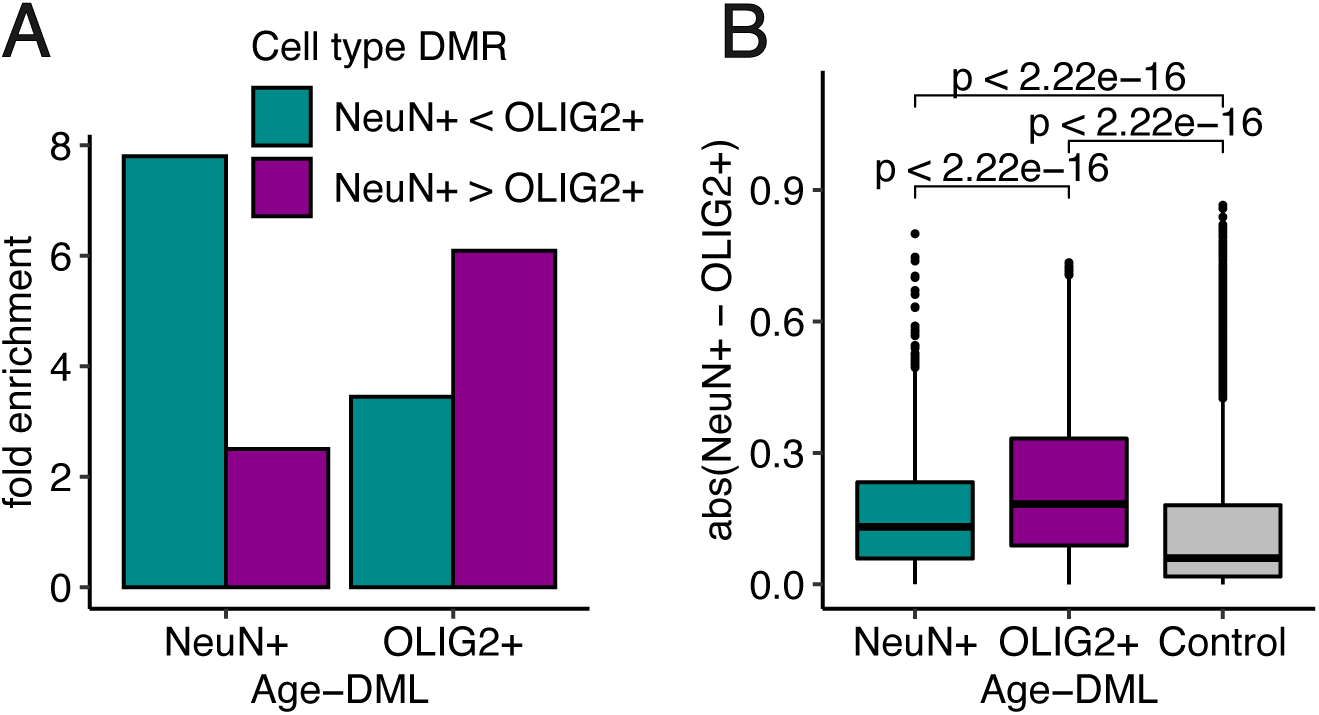
Age-associated DNA methylation changes preferentially occur in cell-type differentiated regions. A) Age-DMLs are highly enriched in cell-type DMRs. Fold enrichment analysis was performed based on the occurrences of age-DMLs in cell type DMRs, compared to random control sets (n=100). B) The absolute CpG methylation difference between cell types is significantly higher for age-DMLs compared to control sets with matched G+C nucleotide contents.

### Dysregulation of cell type identities with aging

Above we have demonstrated that age-associated DNA methylation changes were particularly pronounced for cell-type differentially methylated CpGs. DNA methylation is critically important for regulation of cell type identities [26, 31]. For example, DNA methylation represses expression of cell-type irrelevant genes while promoting expression of cell-type identifying genes [28]. Therefore, we hypothesized that age-associated DNA methylation changes would negatively affect cell-type epigenetic identities. As a representation of epigenetic cell type identities, we measured how different is the configuration of cell type specific DNA methylation. Specifically, for each CpG, we measured the mean methylation levels in NeuN+ and OLIG2+. For example, in a specific CpG, methylation of these two cell types may be 0.2 and 0.9 at a given time. If this site changed its methylation levels to 0.3 and 0.8 at a different age, the epigenetic configuration of this CpG has changed from [0.2, 0.9 in neurons and oligodendrocytes] to [0.3 and 0.8 in neurons and oligodendrocytes]. To depict this difference of epigenetic cell type identity, we calculated the Euclidean distance between these two ‘points’ (in the above example, the distance is 0.14).

The epigenetic distance measured this way exhibited clear patterns. First, epigenetic distance dramatically increased from neonates to toddlers, which was consistent with the developmental shift, also observed in Supplementary Figure 1. Second, while randomly selected non-age-DML CpGs show similar epigenetic distance throughout the age span in the data (Figure 4), sites identified as age-DMLs exhibited clear divergence with age. This pattern was more pronounced for OLIG2+ compared to NeuN+ (Figure 4).

**Figure 4.**
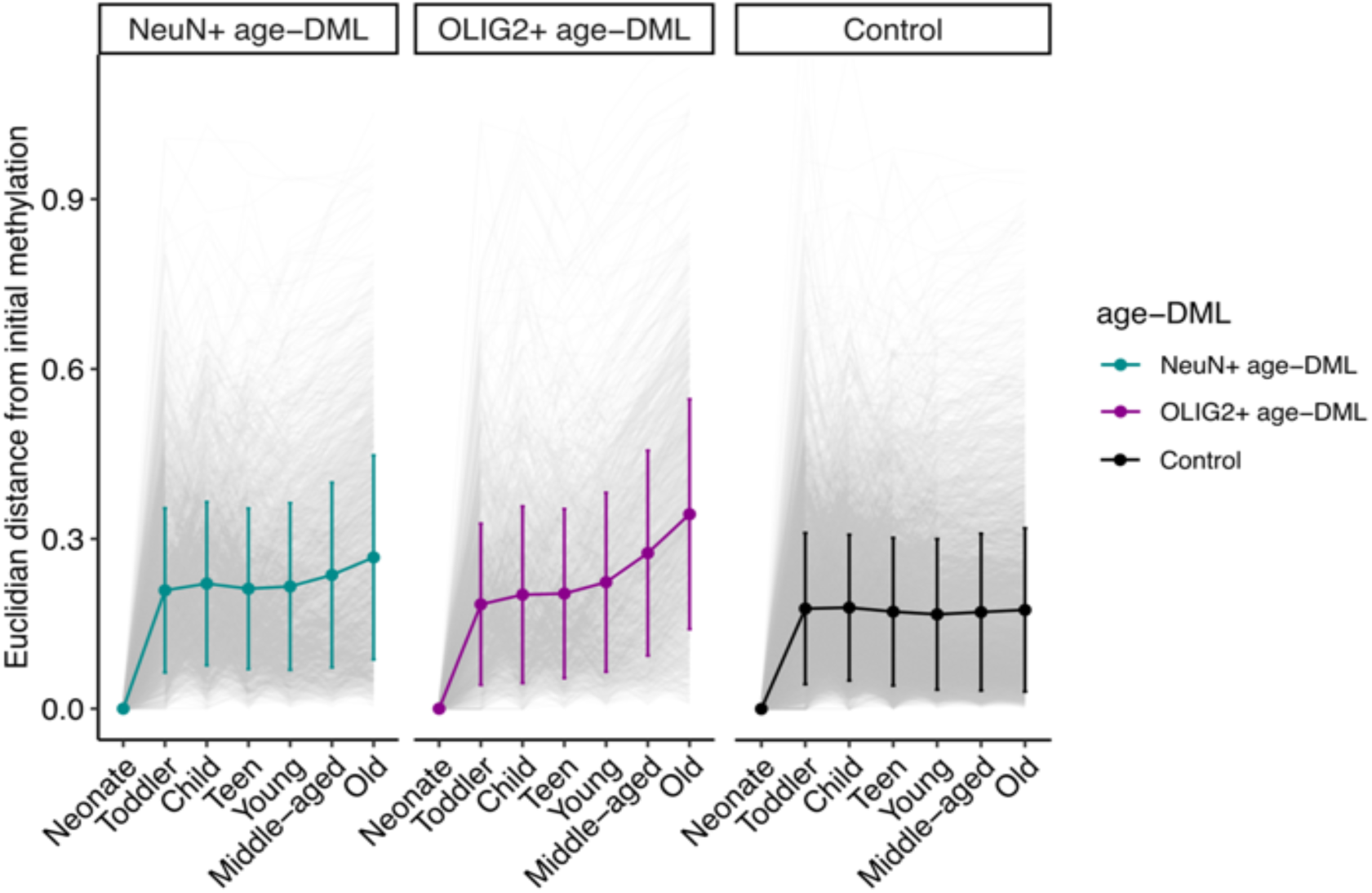
Dysregulation of cell identity with age measured by DNA methylation changes in neurons and oligodendrocytes. Epigenetic distance is calculated by Euclidean distance between points in a two-dimensional plane (DNA methylation of two cell types). Epigenetic distances of cell type identity from the neonate are depicted for different age groups for cell-type specific aging-DMLs. Epigenetic distances of the randomly selected CpG positions are also shown for control.

### Functional Consequences of Age-DMLs

We examined functional implications of age-DMLs. In both cell types, age-hyper DMLs were significantly enriched in CpG islands (Figure 5A). In addition, age-hypo DMLs exhibited strong enrichment in brain enhancers in the dorsolateral prefrontal cortex inferred from Roadmap epigenomics data (Figure 5A). These observations indicate potential impacts on gene expression. As CpG island methylation tends to be negatively correlated with its corresponding gene expression, we examined matched gene expression data and found that genes that accumulate DNA methylation in the promoter (i.e., harboring age-hyper DMLs) experienced decreases of expression with age (Figure 5B). In contrast, genes associated with age-hypo DMLs in promoters tended to increase gene expression with age (Figure 5B).

**Figure 5.**
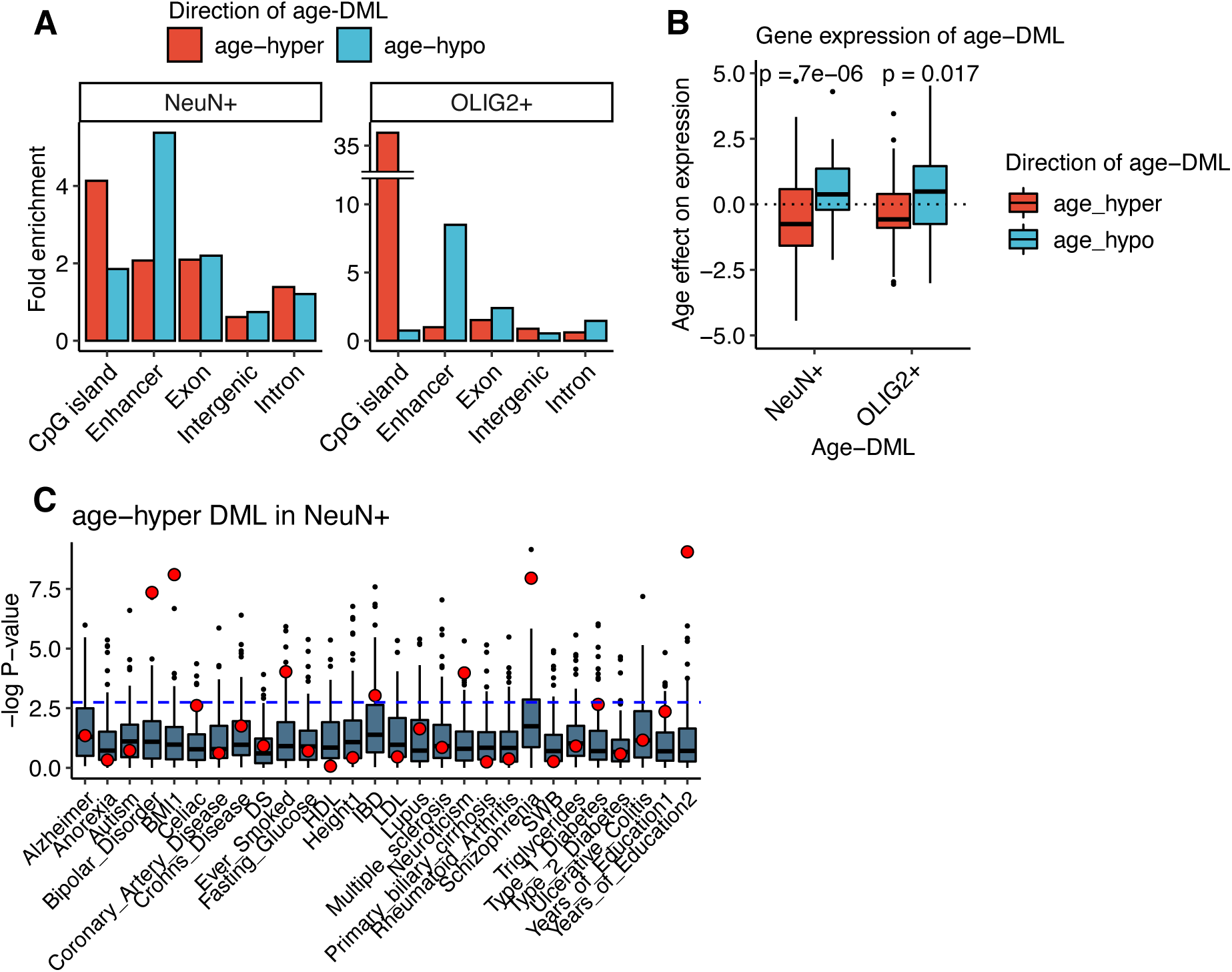
Functional implication and brain trait heritability of age-DMLs. A) Distribution of aging-DML in different functional genomic regions. Aging-hyper DMLs are highly enriched for promoter regions, while aging-hypo DMLs are enriched for brain enhancers. Fold enrichment analysis was performed based on the occurrences of aging-DMLs for each feature compared to random control sets (n=100). B) The effect of age on gene expression for aging-DML using the matched RNA-Seq data (Method). C) Significance levels for genetic heritability in different aging DML and complex traits. Red dots indicate statistical significance of aging-DML for the traits. Boxplots represent results from random control sets (Method).

We further examined the contribution of age-DMLs to genetic heritability associated with human diseases. We performed the stratified linkage disequilibrium score regression [32] to estimate the contribution of age-DMLs to disease and other complex traits using GWAS summary statistics (Figure 5C). Previously, we showed that differentially methylated regions between neurons and oligodendrocytes contribute to neuropsychiatric and neurodegenerative disorders [26]. To avoid a bias derived from the DNA methylation between neurons and oligodendrocytes, we compared the results with two control sets controlling for the cell-type methylation difference and GC ratios (Methods). We found that NeuN+ specific age-hyper DMLs were particularly significantly enriched for several neuropsychiatric disorders as well as body mass index (BMI) and the years of education (Bonferroni adjusted P-value < 0.05, Figure 5C), traits with etiology associated with brain function [32]. NeuN+ hypo-DMLs and OLIG2+ DMLs exhibited fewer enrichments (Supplementary Figure 6).

### Relationship with DNA Methylation Clocks

Recent studies of age predictor models from DNA methylation arrays showed that chronological ages could be accurately predicted based on the DNA methylation of a few hundred CpG positions [20, 22]. Our data, while providing nucleotide-resolution DNA methylation data from the largest number of CpGs possible, lack the adequate sample size to construct a DNA methylation clock. Nevertheless, we examined whether the DNA methylation age predictor could estimate biological ages close to the known chronological ages of our WGBS samples. We applied the Horvath multi-tissue DNA methylation age clock [20], the most well-known age predictor. The Horvath clock accurately predicted methylation ages very similar to chronological ages in our WGBS samples (Figure 6A). The correlation between the estimated DNA methylation ages and the chronological ages were 0.76 and 0.72 for NeuN+ and OLIG2+, respectively, reaffirming that clock CpGs could accurately predict ages from multiple tissues. Given that we have demonstrated that age-DMLs were highly distinct between cell types, we hypothesized that clock CpGs might represent those that show less variability between tissues compared to age-DMLs. Indeed, the multi-tissue age predictor clock CpGs such as Horvath [20] and Levine [22] exhibited reduced DNA methylation variation in 10 tissues we have previously curated (Methods), while age-DML found in neurons and oligodendrocytes showed significantly higher methylation variation compared to clock CpGs as well as to randomly selected CpGs (Figure 6B and Supplementary Figure 7).

**Figure 6.**
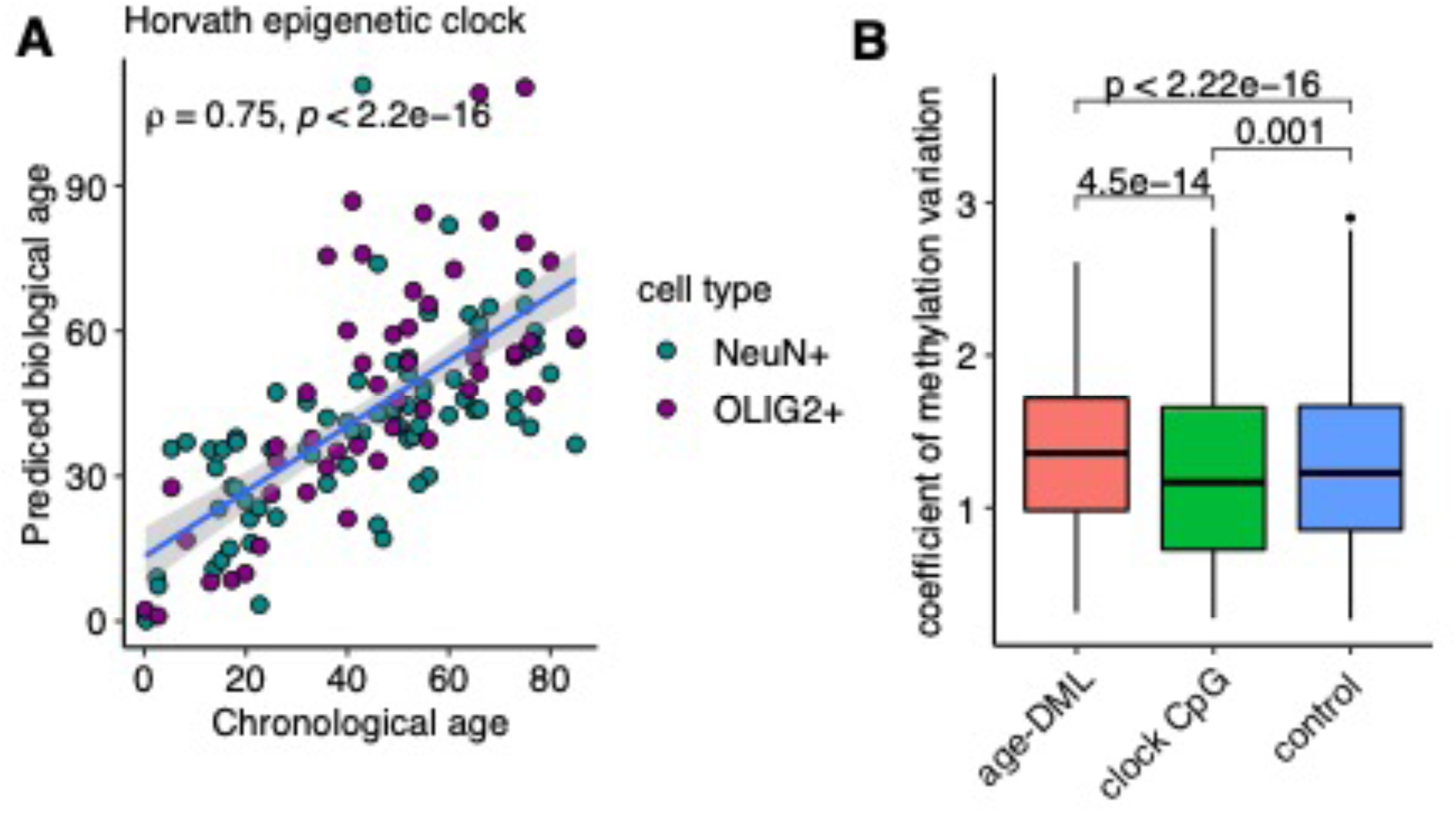
DNA methylation variation of multiple tissues in clock CpGs. A) Horvath estimated DNA methylation age is highly correlated with chronological age of WGBS samples. B) Coefficient of methylation variation resulting from 10 different WGBS tissues shows highly variable DNA methylation for age-DML and reduced DNA methylation variation for clock CpGs.

## DISCUSSION

Our study is, as far as we are aware, the most comprehensive study of age-associated DNA methylation changes at the whole-genome scale, by examining 23.6 million CpGs, and from distinctive cell populations. This work emphasizes the importance of employing unbiased whole-genome profiling of DNA methylation with cell-type resolution to obtain a comprehensive landscape of age-associated methylation variation. Across the two different data sets, aging explained more than 10% of total variation in DNA methylation, which is greater than the estimated effect of sex in both data sets, and the effect of schizophrenia diagnosis in Mendizabal et al.’s data.

We first show that the majority of age-associated variation in DNA methylation can be explained by the effect of DNA methylation in early ages. Using DNA methylation levels at neonates as a proxy, nearly 8.1% of total variation in genomic DNA methylation could be explained by the level of DNA methylation in the early stage (Figure 2). The term ‘epigenetic drift’ has been used in the context of age-associated DNA methylation changes [12–14]. The term drift itself explicitly refers to random changes, and epigenetic drift implicates stochastic change of DNA methylation. However, having demonstrated that the direction of DNA methylation change is significantly related to the DNA methylation level in early age, our study shows that CpG positions that were lowly methylated in neonates tended to increase methylation with age, while those with heavy initial methylation tended to lose DNA methylation. One hypothesis is that epigenetic drift occurs due to the noise introduced by the imperfect DNA methylation machinery accumulated in every cell cycle (e.g., [12]). We show here that neurons show age-associated DNA methylation changes even though they are largely post-mitotic. Therefore, epigenetic drift may not necessarily be associated with cell division. Epigenetic drift may instead be associated with deterioration of other maintenance mechanisms of the epigenomes.

Even though the overall trend of a clear relationship between DNA methylation levels and age-associated DNA methylation change is consistent in the two cell populations, the specific CpGs that show age-associated methylation changes are highly distinct between cell types. Moreover, we show that CpGs that distinguishes cell types are particularly prone to age-associated DNA methylation changes. Recent studies, including those from our group, have shown that DNA methylation and other epigenetic landscapes of different cell types are significantly divergent, and those divergently methylated positions are enriched in genomic regions implicated in diseases (e.g., [26, 31, 33]). We have demonstrated that specific DNA methylation landscapes of distinct cell types become perturbed with aging, as a consequence of epigenetic drift and the divergent DNA methylation landscapes of different cell types. In other words, aging is associated with dysregulation of cell type specific epigenetic identities.

An exciting recent development in the aging research is the identification of Aging CpG clocks, which are subsets of CpGs that can predict the biological and phenotypic ages with high accuracy. How can we reconcile aging CpG clocks with our findings that age-associated DNA methylation changes occur in a highly cell type specific manner? It should be noted that our study and aging CpG clocks use fundamentally different approaches. The goal of aging CpG clock studies is to identify predictors of aging from the CpGs that are present in large studies, which are typically those included in widely used DNA methylation arrays. The specific CpGs within the clocks often do not show higher correlation with age than other CpGs [20]. Rather, these two types of CpGs represent two different aspects of age-associated changes of DNA methylation. Nevertheless, interestingly, clock CpGs appear to have less variability of DNA methylation between tissues compared to randomly selected CpGs as well as age-DMLs. Among the clock CpGs, those selected from a single tissue (Hannum clock, [9]) show a similar level of variability with the cell-type age-DMLs (Supp Figure 8). Results of our studies indicate that dysregulation of epigenetic cell-type identities and current DNA epigenetic clocks may represent different aspects of genomic and cellular age-associated DNA methylation alterations. Our study emphasize future needs to integrate these distinctive features of epigenetic aging to aid comprehensive understanding of molecular processes of aging.

## Methods

### Whole-genome bisulfite sequencing data processing

To investigate comprehensive brain cell-type DNA methylation changes with aging, we used our previously published WGBS data set of 53 NeuN+ and 42 OLIG2+, which comprised samples from individuals of different ages (range from 25 yrs to 85 yrs) [26]. We also collected brain cell-type WGBS data from neonates, toddlers, and teens to examine DNA methylation trajectories across different age groups and infer the initial cell-type DNA methylation state.

WGBS reads were quality trimmed with Cutadapt (ver. 3.1) using a default setting [34]. The trimmed reads were mapped to the human reference genome (hg19) using Bismark (ver. 0.23) with Bowtie 2 mode [35]. Duplicated reads were further processed and filtered by the deduplicate module in Bismark (deduplicate_bismark). We removed lowly mapped CpG positions (average mapped reads less than 5).

### Identification of CpG positions with extreme DNA methylation changes with aging

For each CpG position, we fitted a generalized linear model using the arcsine link function to estimate the age effect. The model fitting is conducted utilizing the DSS (ver. 2.4) Bioconductor package [29]. We considered bisulfite conversion rates, sex, postmortem intervals, brain bank, and disease status as covariates. A hypothesis test of the age effect for each CpG site was performed using the Wald test using the estimated coefficient and standard error from the fitted model. False discovery rate (FDR) is computed using the Benjamini-Hochberg method. The coefficients of age from the DSS analyses are estimated from the GLM framework, making it difficult to interpret the explicit meaning of methylation differences compared across CpG sites. Thus, we also fitted general linear models using fractional methylation to estimate biologically interpretable methylation level differences with aging.

### RNA-Seq data processing

RNA-Seq data from the matched with WGBS samples were collected from previous studies [26, 27]. Raw sequencing reads were quality trimmed with Trimmomatic (ver. 0.39) [36] and then mapped to the human reference genome (hg19) using STAR (ver. 2.7) with the following options: --alignSJDBoverhangMin 1 --outFilterMismatchNmax 3 --outFilterMultimapNmax 10 -- alignSJoverhangMin 10 --twopassMode Basic. We removed any secondary alignments and duplicated reads using Samtools (ver. 1.13) to ensure that only uniquely mapped reads were retained for further analyses [37]. We calculated the gene expression using htseq (ver. 0.11.2) using intersection-strict mode by the exonic regions [38]. We quantified protein-coding genes using the human Ensembl annotation (GRCh37.87).

### Disease heritability using stratified LD score regression

To measure the contribution of age-DML to the genetic risk of disease and complex traits, we performed the stratified LD score regression analysis. We followed the same processing steps and used the list of GWAS traits described in the previous work [30]. Briefly, we examined extended genomic regions of age-DML (25 kbp on both sides of focal age-DML) to improve the confidence intervals of the estimates. As a statistical control, we performed partitioned stratified LD score analyses using CpG positions that are differentially methylated between neurons and oligodendrocytes. G+C nucleotide contents were matched based on the GC ratio of 1kbp window (+-500bp of age-DML).

### Variation of DNA methylation across tissues

To estimate the degrees of DNA methylation variation across different tissues, we used the processed WGBS data from 9 tissues (placenta, sperm, hair follicle, adrenal gland, liver, colon, ovary, embryonic stem cell, and b-cell) [39]. To measure the variation of DNA methylation, we used the modified coefficient of variation proposed by Chatterjee et al. [40], which corrects for the dependence on the mean value as follows:,

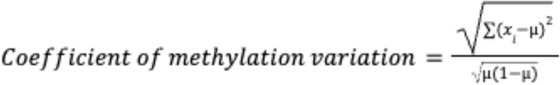, where *x_i_* denotes fractional methylation of i-th tissue (n is the number of tissues in the data) and *μ* denotes the mean methylation value.

## Acknowledgments

This study is partially supported by NSF (EF-2021635), NIH (HG011641), and ICB (27KK01) grants to SVY and a CRIS Contra El Cancer Foundation (PR_TPD_2020-19) to IM.

