## Supplementary Figures for "Human Brain Aging is Associated with Dysregulation of Cell-Type Epigenetic Identity"

**Figure S1.** Principal component analysis of whole-genome CpG methylation values. The first principal component (PC1) which explains 10.3% of a total variation of DNA methylation distinguishes the age groups.

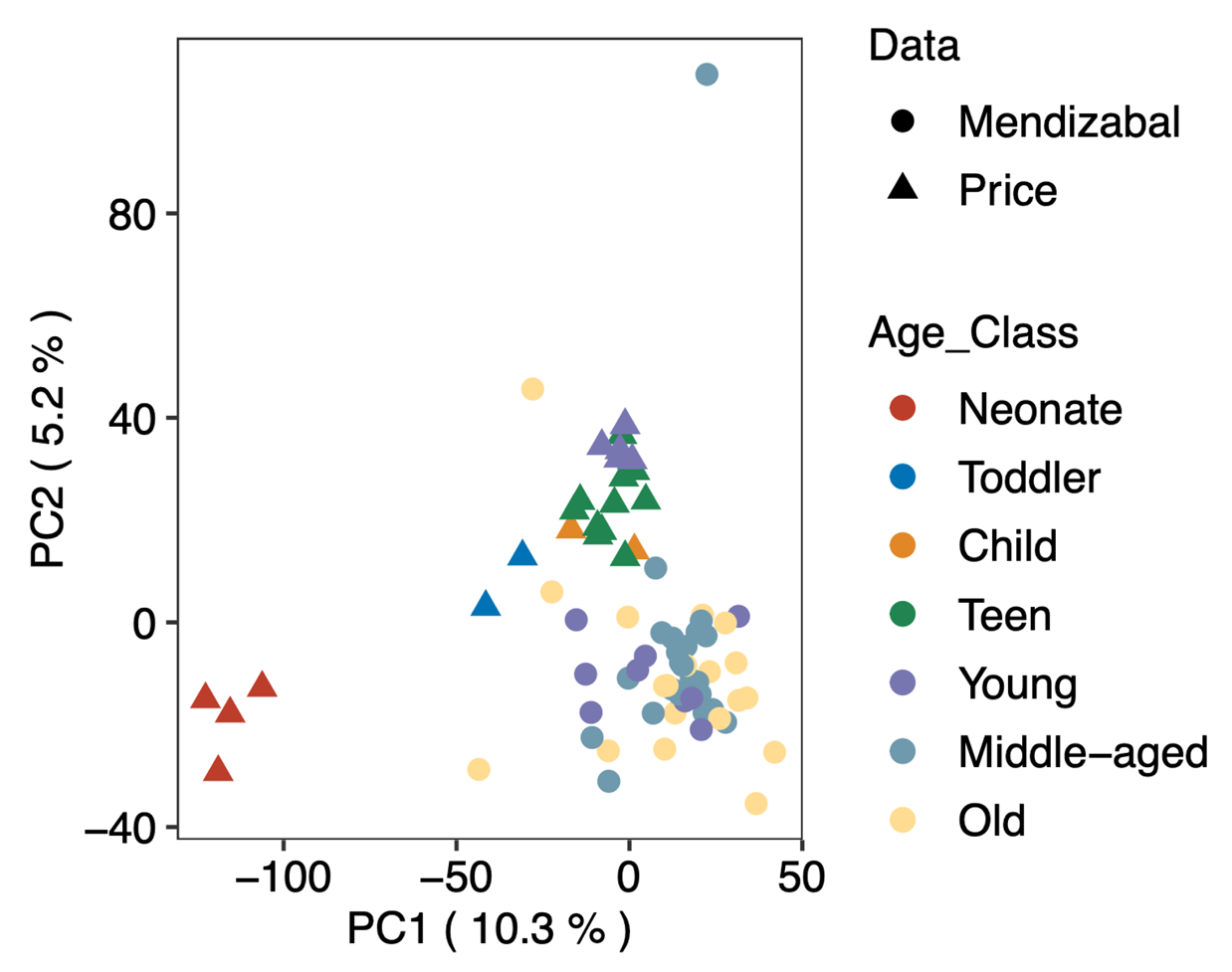

**Figure S2.** A relationship between mean methylation and the effect of age on methylation. For each CpG site, we fitted a linear model to estimate the age effect on DNA methylation adjusted for other biological variables (post-mortem interval, sex, disease status, and bisulfite conversion rate). For an illustration purpose, only 10,000 randomly selected CpG sites were displayed in the plots. (A) Y-axis indicates mean methylation levels of neonates for the corresponding CpG site. (B) Y-axis indicates mean methylation levels of samples with age < 20 for the corresponding CpG site.

**A**

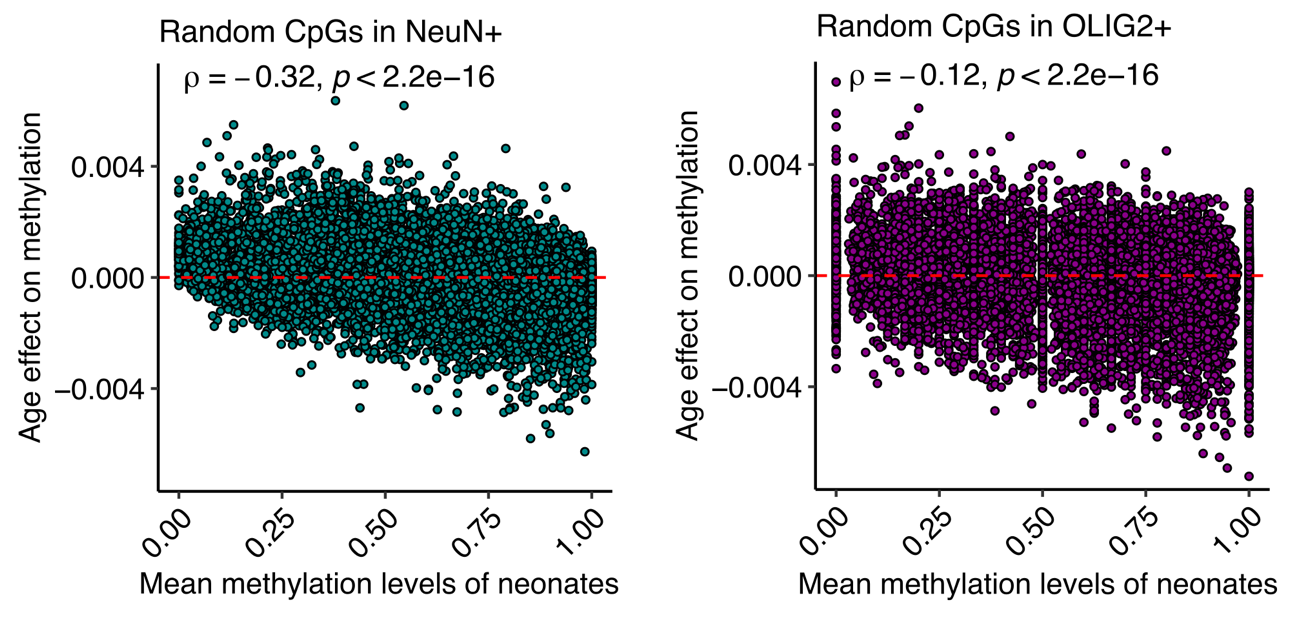

**B**

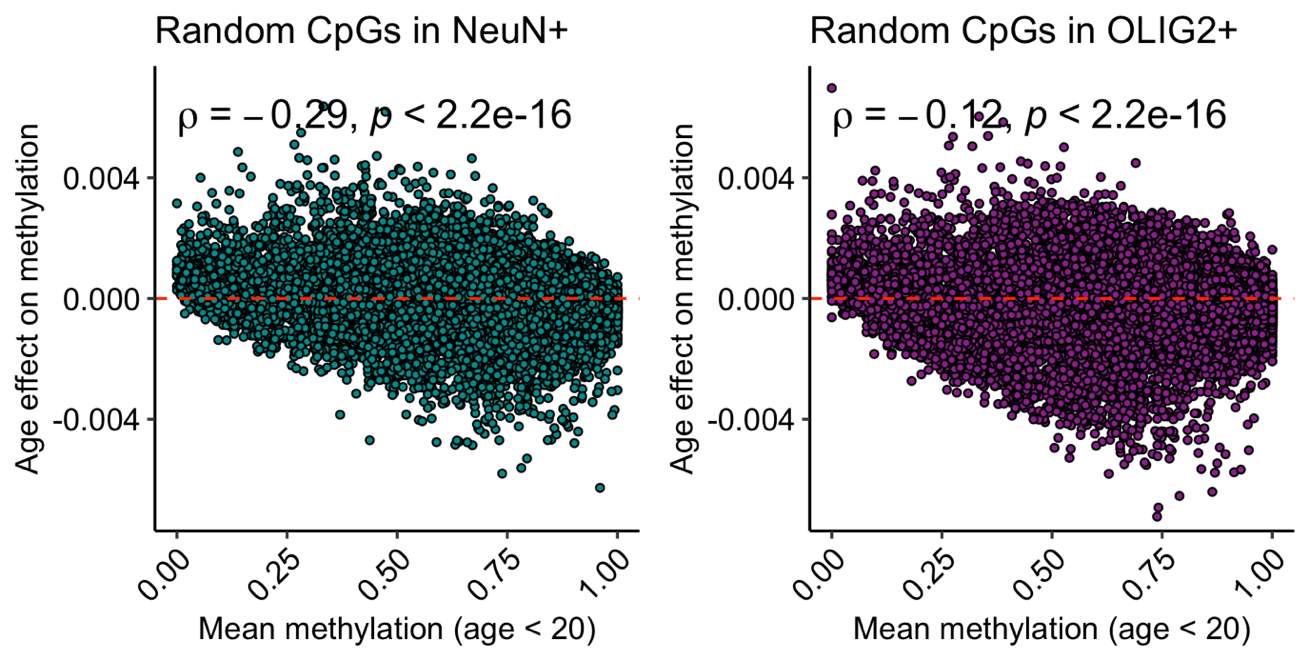

**Figure S3.** Distributions of whole-genome CpG methylation across age groups. NeuN+ samples were ordered by age (from bottom to top). For the illustration purpose, CpG sites with fractional methylation greater than 0.9 or less than 0.1 for more than 90% of samples were excluded. Also, samples with low bisulfite conversion rate (<95%) were excluded. Sample index is denoted as ‘sample name_chronological age’

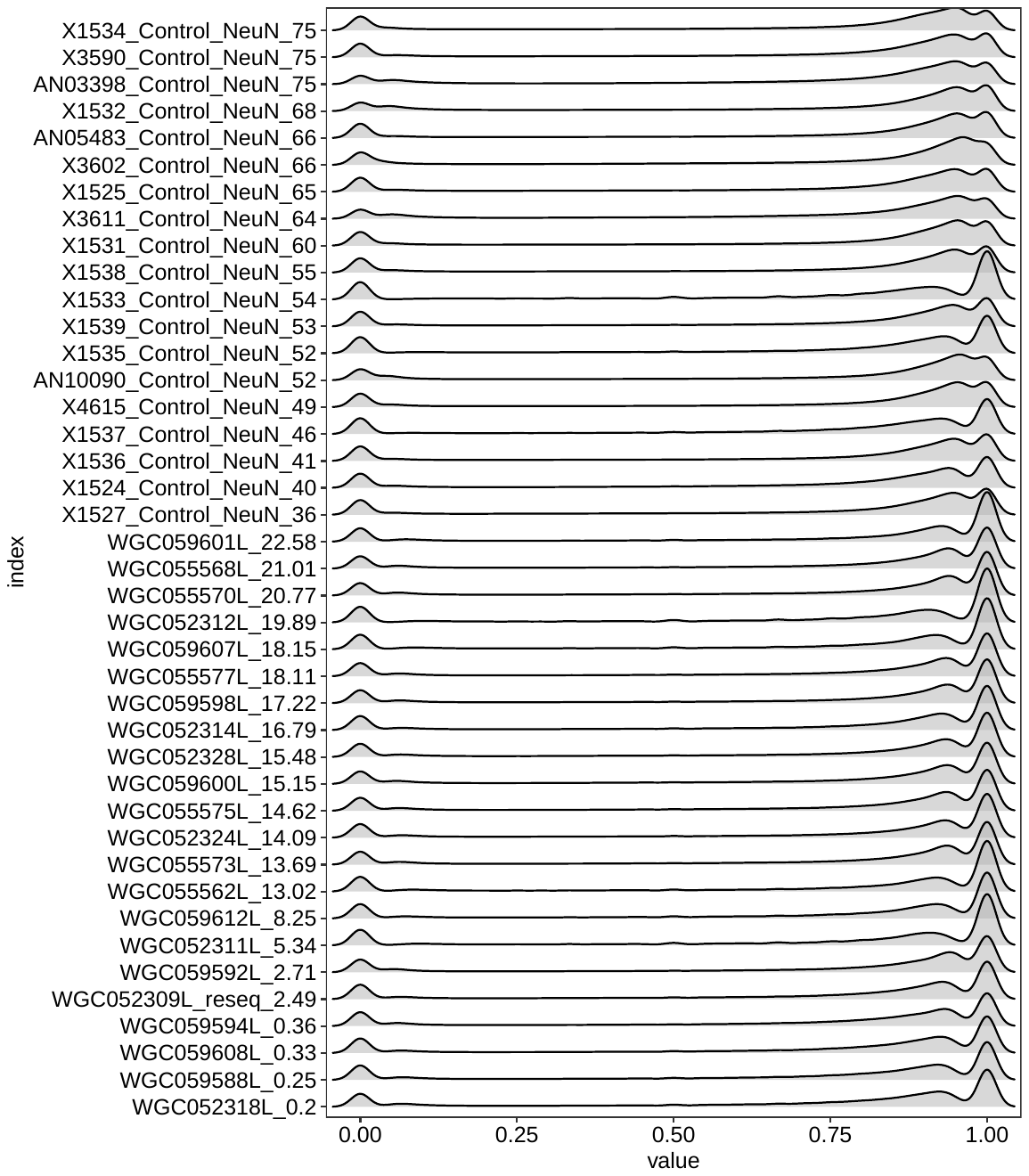

**Figure S4.** Concordance of DSS statistics between results from the combined data set (Mendizabal and Price) and results from using only Mendizabal data set. CpG sites with disconcordant DSS statistics from two analyses were colored in red.

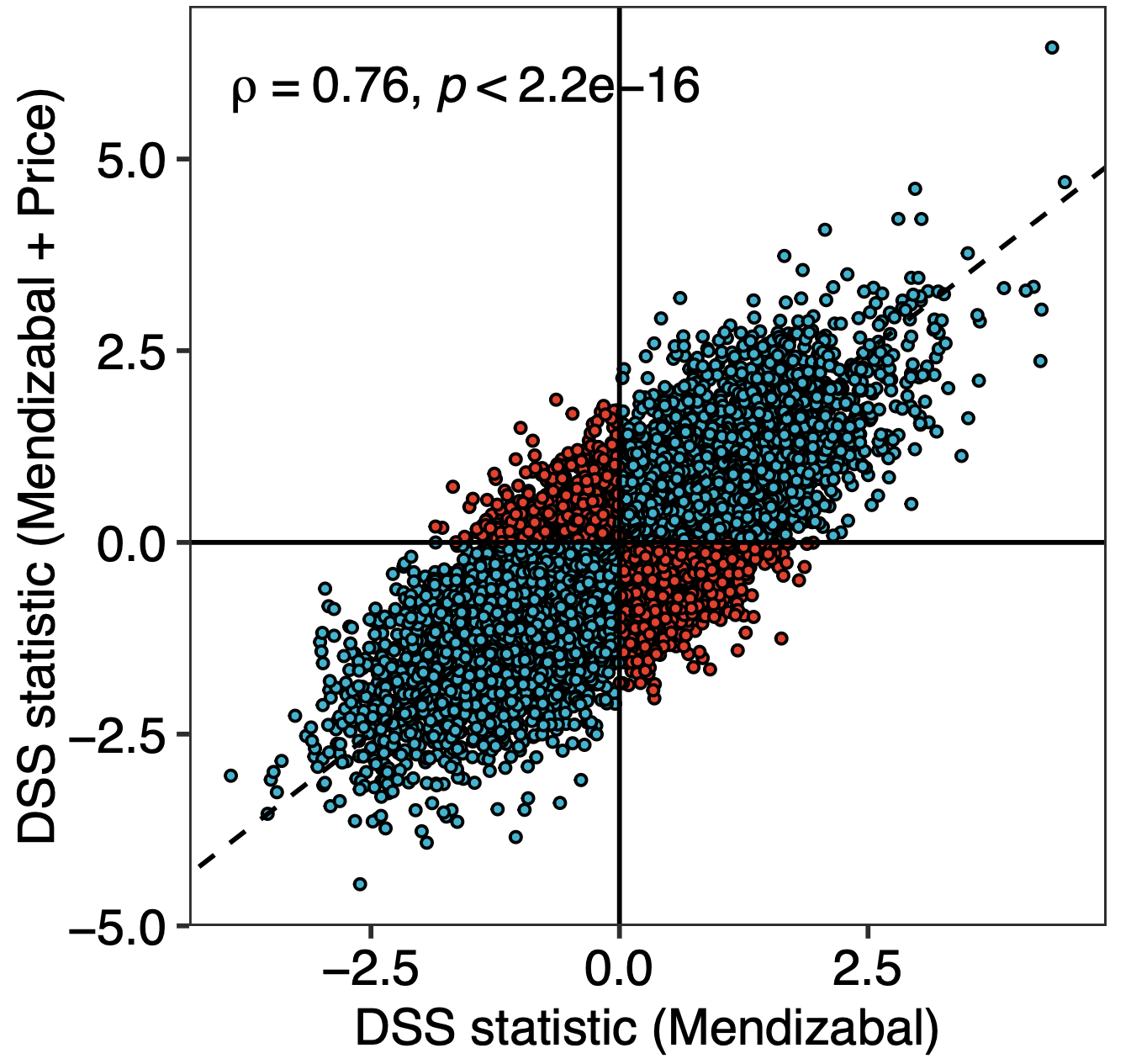

**Figure S5.** Trajectories of cell type methylation difference with age. Relative methylation difference between NeuN+ and OLIG2+ was calculated for the samples collected from the same brain tissue.

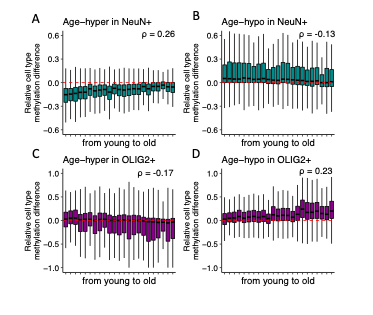

**Figure S6.** Significance levels for genetic heritability in different aging DML and complex traits. Red dots indicate statistical significance of aging-DML for the traits. Boxplots represent results from random control sets (Method).

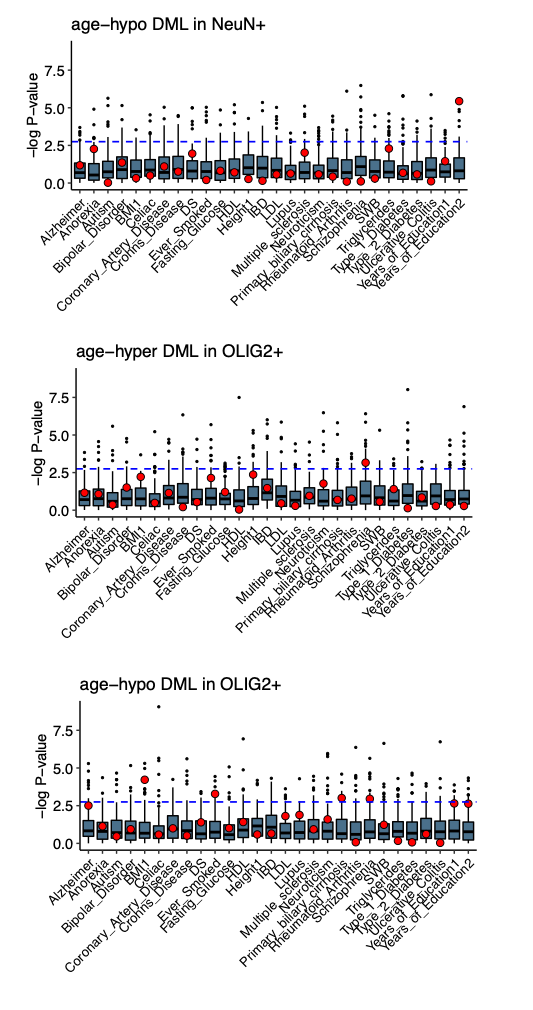

**Figure S7.** DNA methylation variation of multiple tissues in clock CpGs. Coefficient of methylation variation resulting from 10 different WGBS tissues shows highly variable DNA methylation for age-DML and reduced DNA methylation variation for clock CpGs.

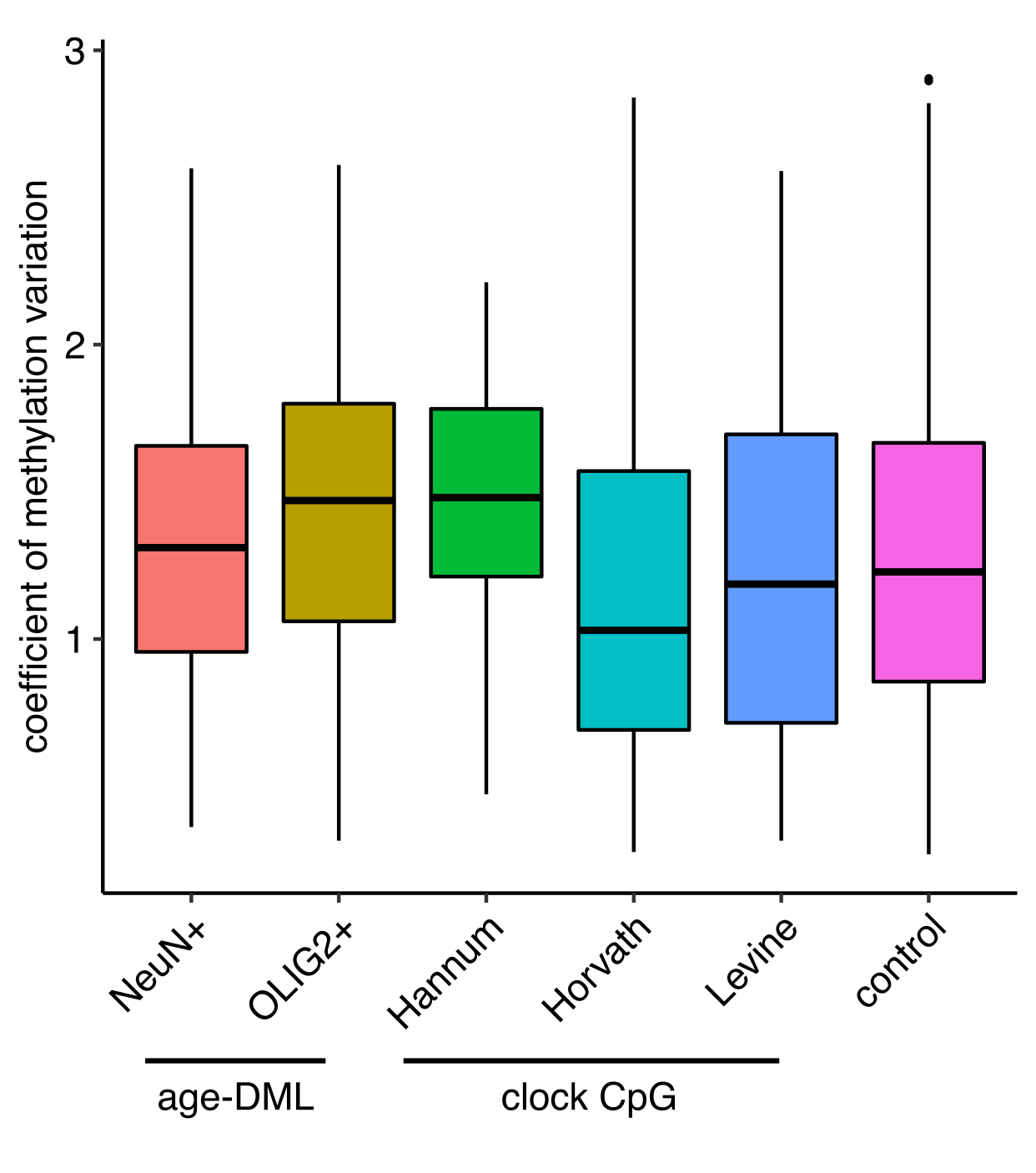
